# ESMRank reveals a transferable axis of protein mutational constraint from overlapping variant effect assays

**DOI:** 10.64898/2026.02.26.708185

**Authors:** Riccardo Arnese, Gennaro Gambardella

## Abstract

Proteome-wide interpretation of missense variation is constrained not only by predictive model performance but also by the absence of principled methods to reconcile heterogeneous multiplexed assays of variant effect (MAVEs) into a unified representation of mutational constraint. We show that redundancy among partially overlapping deep mutational scanning experiments encodes a reproducible ordinal signal that can be recovered despite differences in assay scale and readout. We introduce variant soundness, an overlap-aware framework that aligns within-assay rankings and aggregates them across experiments to derive an assay-agnostic, within-protein measure of mutational tolerance. Applying this approach to about 1,100 MAVEdb score sets spanning >2M variants reveals a coherent constraint landscape enriched for structural stability determinants, including residue burial, packing perturbation magnitude, and domain architecture. By aligning learning objectives with this intrinsic ordering, we develop ESMRank, a sequence-based learning-to-rank predictor integrating protein language model representations with physicochemical descriptors. Under strict protein-level partitioning, ESMRank outperforms widely used stability and fitness predictors across the Human Domainome, ProteinGym stability assays, and VariBench folding kinetics. Without clinical supervision, the reconstructed constraint axis is enriched for ClinVar pathogenic variants and stratifies genes by mechanistic disease classes. In CFTR, predicted constraint tracks folding efficiency, channel activity, and pharmacological rescue. These findings establish experimental overlap as a scalable resource for extracting transferable mutational ordering and for building mechanistically interpretable, proteome-wide variant effect predictors.

## INTRODUCTION

Missense variants are the most prevalent form of protein-coding genetic variation in humans, with each individual carrying thousands of amino acid–altering substitutions (1). Although a minority are pathogenic, most remain uncharacterized, and only a small fraction of the millions of possible missense variants have been experimentally or clinically annotated (2). This gap underscores the need for scalable approaches to variant interpretation that generalize across proteins and biological contexts.

Multiplexed assays of variant effect (MAVEs), including deep mutational scanning, can quantify functional consequences for thousands to millions of variants per protein (3–6). These studies have yielded detailed mutational landscapes that connect sequence changes to biophysical and structural constraints, including residue burial, packing, and interaction interfaces (5,6). However, MAVEs are intrinsically heterogeneous (5,6). Assays differ in readout, design, dynamic range, cellular context, and scoring conventions, so effect magnitudes are often not directly comparable across experiments, even when they test overlapping variants (5–8). As MAVE datasets accumulate, partial redundancy between studies is increasingly common, yet overlap is typically handled ad hoc (9) or treated as noise rather than leveraged as an explicit source of cross-assay information (3,4,9). Consequently, mutational landscapes remain fragmented, limiting the extraction of transferable constraint signals from the experimental record.

At the same time, computational prediction has advanced rapidly. Protein language models and structure-aware predictors can learn strong priors from large-scale sequence and structural data and estimate mutational effects in zero-shot settings (10,11). A natural next step is to improve these models using functional measurements, but naïvely fine-tuning on limited or pooled MAVEs often yields limited gains and can reduce generalization (12), in part also because heterogeneity confounds direct regression across assays (13,14). Importantly, while absolute effect sizes vary substantially across experimental contexts, the relative ordering of variant effects within a protein is often more reproducible (14,15). This suggests that ordinal structure, rather than scale, may constitute the more stable and transferable signal, particularly for variants that disrupt folding (16), a frequent upstream bottleneck impacting diverse downstream readouts.

Here we address these challenges through an overlap-aware integration framework that uses partially overlapping MAVEs to recover reproducible mutational structure from heterogeneous experiments. We quantify cross-assay rank consistency on shared variants and derive a consensus variant score (here termed “variant soundness”) (17) that emphasizes ordinal agreement while reducing sensitivity to assay-specific scaling. Applying this framework to over two million variants across more than one thousand experiments yields coherent gradients of constraint aligned with structural environment and domain organization.

Recognizing that this integrated signal is inherently relative within proteins, we then formulate variant effect prediction as a learning-to-rank problem (18,19) and develop ESMRank, a sequence-based model that integrates protein language model embeddings with complementary physicochemical descriptors to capture both evolutionary and biophysical determinants of variant effect. Under strict protein-level partitioning and independent benchmarking, we evaluate its performance on stability- and abundance-focused datasets and examine how the reconstructed constraint landscape relates to clinical pathogenicity and disease mechanism. As a case study, we further assess whether predicted mutational tolerance in CFTR aligns with experimentally measured folding efficiency, channel function, and pharmacological responsiveness.

Our results establish experimental overlap as a scalable statistical resource for harmonizing heterogeneous MAVEs and show that aligning both integration and modeling objectives with ordinal structure enables transferable and mechanistically interpretable maps of missense variants across proteins.

## MAIN

### 1. Overlap-aware integration of MAVEs yields a unified mutational signal

Multiplexed assays of variant effect (MAVEs) have generated extensive functional measurements across proteins, organisms, and assay modalities. However, heterogeneity in dynamic range, experimental context, and coverage limits direct comparability and obscures global patterns of constraint. The MAVEdb release analyzed here comprises 1,122 score sets spanning 2,176,052 mutations across 596 proteins, predominantly human (Fig. 1A). Notably, more than half of these datasets (581 score sets; 1,143,050 mutations) partially or completely overlap in the variants assayed (Fig. 1B), forming a dense yet heterogeneous web of redundant measurements.

**Figure 1.**
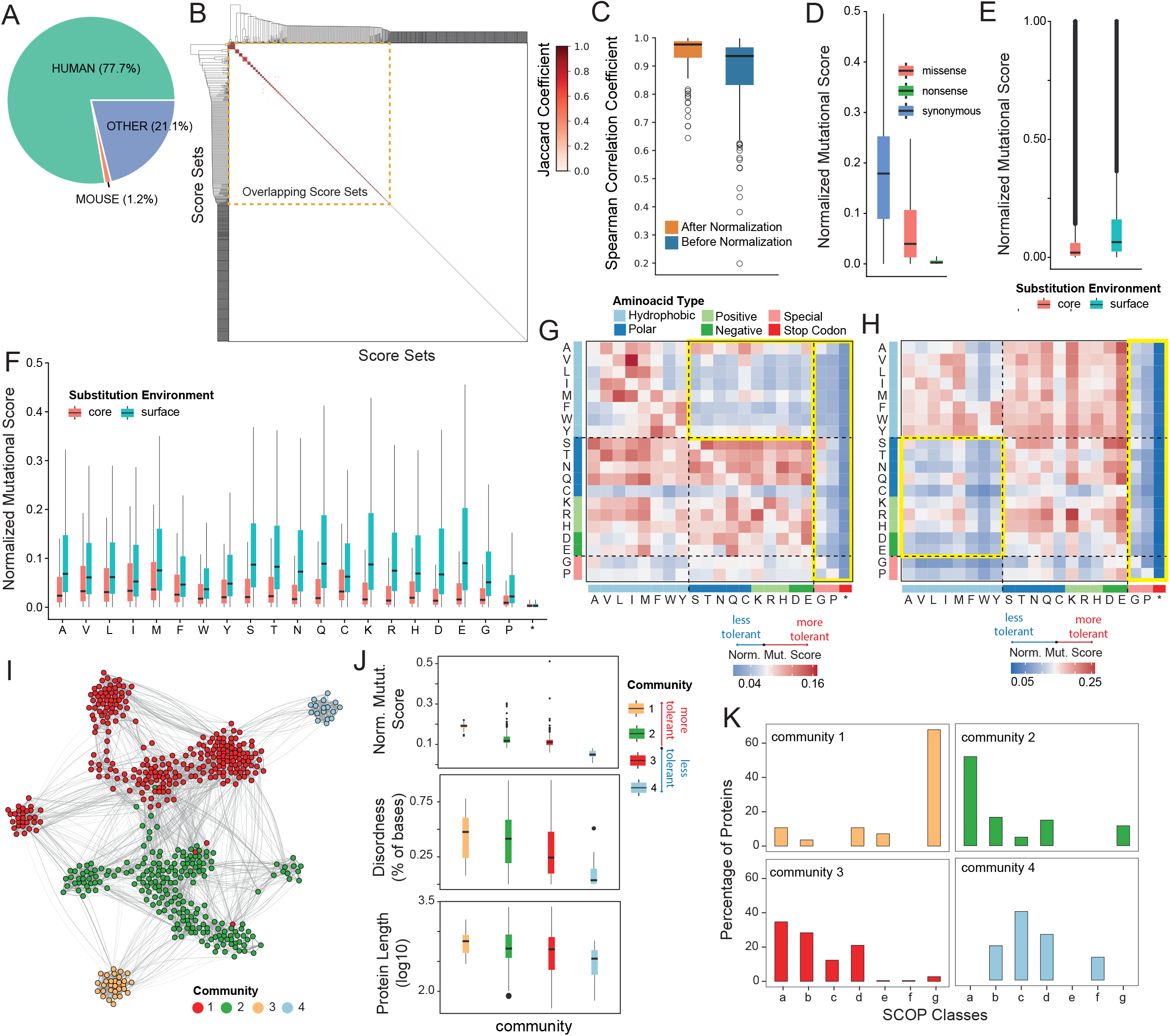
Overlap-aware integration of MAVEs yields a unified and biologically coherent mutational signal. (**A**) Species composition of the MAVEdb score sets analysed in this study, showing a predominance of human proteins. (**B**) Pairwise overlap among MAVEdb score sets quantified using the Jaccard coefficient. A large block of partially or fully overlapping experiments (dashed orange box) highlights redundancy across independent assays for the same proteins. (**C**) Cross-experiment coherence before and after integration. Spearman correlation coefficient between individual DMS score sets is substantially increased following soundness normalization, indicating effective suppression of assay-specific noise. (**D**) Distribution of normalized mutational scores for synonymous, missense, and nonsense variants after normalization, showing enhanced separation of functional classes. (**E**) Comparison of normalized mutational scores for substitutions occurring in buried (core) versus surface-exposed residues showing globally reduced tolerance for substitutions in protein cores relative to surfaces. (**F**) Amino-acid–specific mutational tolerance stratified by structural environment, showing globally reduced tolerance for substitutions in protein cores relative to surfaces with nonsense variants being uniformly deleterious. (**G**) Environment-specific substitution matrix for buried residues. Hydrophobic-to-polar or charged substitutions in the protein core are strongly deleterious, reflecting classical stability constraints. (**H**) Environment-specific substitution matrix for surface residues, revealing distinct electrostatic and functional sensitivities, with many charged and polar substitutions disproportionately deleterious at exposed positions, consistent with disruption of interaction interfaces and solvent-mediated networks. (**I**) Mutational similarity network constructed by representing each protein through its combined core and surface substitution signatures. Nodes represent proteins and edges connect proteins with similar mutational response profiles; colours indicate network communities. (**J**) Protein-level properties across network communities. Communities ordered by average mutational tolerance show systematic differences in normalized mutational score, intrinsic disorder content, and protein length, with the most tolerant community enriched for longer and more disordered proteins. (**K**) SCOP class composition of network communities. The most tolerant community (community 3) is enriched for SCOP class g domains, including small, often metal-binding modules such as zinc fingers, whereas the least tolerant community (community 4) is enriched for β-rich folds (SCOP classes b–d), indicating that mutational tolerance reflects the interplay between global sequence properties and local domain architecture rather than global fold alone.

While assay-specific noise precludes naïve aggregation, overlapping experiments provide independent measurements of shared variants that can be leveraged to extract a consensus signal. We therefore developed variant soundness, an overlap-aware metric that quantifies the consistency with which a variant is ranked across assays using rank alignment and Reciprocal Rank Fusion (17) (RRF, Methods). Soundness profiles exhibited increased cross-experiment coherence (Fig. 1C), correlating more strongly with their constituent Deep Mutational Scanning (DMS) assays than those assays correlated with one another, indicating suppression of assay-specific noise. To enable joint analysis across the full dataset, RRF-integrated profiles were then combined with non-overlapping score sets and normalized onto a common scale (Methods), maintaining separation of synonymous, missense, and nonsense mutations (Fig. 1D), thus yielding an assay-agnostic representation of variant effects. A simple metadata-based survey using keyword matching indicated that stability- or abundance-associated readouts constitute a substantial fraction of score sets (Fig. S1). These assays, however, span diverse experimental implementations and cellular contexts (e.g., mainly mammalian and yeast cells) and are interspersed with binding, activity, and composite fitness measurements (3,4).

Analysis of the integrated dataset revealed pronounced biophysical structure. Buried residues were substantially less tolerant than surface-exposed positions (Fig. 1E–F), whereas nonsense mutations were uniformly deleterious across environments (Fig. 1F) (20). Substitution matrices delineated distinct constraint regimes: hydrophobic-to-polar or charged substitutions were strongly deleterious in protein cores (21,22), consistent with stability constraints, whereas surface residues displayed selective sensitivity to charged and polar substitutions, consistent with disruption of intermolecular interfaces (Fig. 1G–H) (21,22). To test whether this organization reflects graded biophysical costs, we jointly stratified variants by solvent accessibility and side-chain volume perturbation (|ΔVolume|). Tolerance increased with exposure and decreased with perturbation magnitude (Fig. S2), with large volume changes markedly more deleterious at buried residues but comparatively attenuated at exposed positions. A linear mixed-effects model confirmed a strong interaction between exposure and |ΔVolume| (t statistic = −16.05), demonstrating that the impact of packing perturbation scales with residue burial. These results indicate that the integrated axis encodes exposure-dependent structural constraints consistent with stability-related packing effects, alongside distinct sensitivities at exposed sites. The prominence of these stability-associated gradients across heterogeneous assays is consistent with the notion that folding defects represent a shared bottleneck across diverse functional readouts (16).

Next, to examine whether these residue-level trends extend to higher-order organization, we summarized each protein by its combined core and surface substitution signatures and constructed a mutational similarity network that groups proteins by shared patterns of mutational response (Fig. 1I; Methods). Ordering network communities by average tolerance (Fig. 1J, upper panel) revealed systematic associations with global sequence properties, with the most tolerant community comprising longer proteins (Fig. 1J, lower panel) enriched in intrinsically disordered regions (Fig. 1J, middle panel) (23–26). This organization extended to the domain level, as SCOP class enrichment analysis revealed distinct compositional biases across communities (Fig. 1K). The most tolerant community (community 1) was enriched for SCOP class g domains, including small, often metal-binding modules such as zinc fingers, consistent with architectures in which compact, locally constrained domains are embedded within long and disordered sequences (27). In contrast, the least tolerant community (community 4) was enriched for β-rich folds and more compact, structurally constrained architectures (SCOP classes b–d) (27). These observations indicate that mutational tolerance is consistent with the interplay between global sequence properties and local domain architecture rather than global fold alone (23–27).

Finally, to assess clinical relevance, we compared integrated scores with ClinVar missense variant annotations (2). Following intersection at the protein level, 1,069 pathogenic or likely pathogenic variants and 310 benign or likely benign variants were represented within the MAVEdb-integrated dataset. Pathogenic variants were strongly enriched at the deleterious end of the integrated axis relative to benign variants (Wilcoxon rank-sum test, P = 2.0 × 10□^12^; Fig. S3). The cumulative distribution of scores revealed a pronounced leftward shift for pathogenic variants, consistent with the predominance of loss-of-function mechanisms in human genetic disease. These results indicate that the unified mutational landscape captures biologically meaningful constraint patterns that extend to human pathogenic variation.

Together, these results demonstrate that overlap-aware integration of heterogeneous MAVEs reveals a coherent and biologically structured axis of mutational constraint. This unified landscape captures both local packing sensitivities, higher-order architectural influences, and patterns of human pathogenic variation while remaining agnostic to assay-specific scale. By distilling diverse experimental measurements into a shared constraint framework, it enables principled prioritization of missense variants across diverse proteins and experimental contexts.

### 2. ESMRank: A learning-to-rank AI framework for protein variant effects prediction

Because the integrated signal is inherently ordinal, reflecting relative mutational tolerance within proteins rather than absolute effect magnitudes, we formulated variant effect prediction as a learning-to-rank problem (18,19,28,29). Unlike regression models that attempt to reproduce heterogeneous score scales, a ranking formulation directly optimizes the relative ordering of variants within each protein, aligning naturally with the consensus structure recovered by variant soundness.

We developed ESMRank, a sequence-based model trained on 1,033,002 soundness-normalized variants using a protein-level stratified cross-validation scheme to prevent information leakage across proteins and experiments (Methods). The model employs LambdaMART (30), a gradient-boosted decision tree implementation of pairwise learning-to-rank, to optimize discrimination between more and less deleterious substitutions within each protein.

Given a protein sequence, ESMRank outputs a ranked list of all possible single-amino-acid substitutions, placing predicted loss-of-function variants at the bottom and neutral or function-enhancing substitutions toward the top (Fig. 2A). Variants are encoded using a multimodal feature representation that integrates deep embeddings from the ESM-2 protein language model (31) with a curated set of shallow biophysical, structural, and positional descriptors for a total of 18 descriptors (Supplementary Table 01). Deep features, derived from ESM2, capture global sequence context and sequence-encoded structural perturbations, including embedding-space differences, attention-derived residue–residue contact information, and masked marginal residue-probability shifts, whereas shallow features summarize local physicochemical properties (Methods).

**Figure 2.**
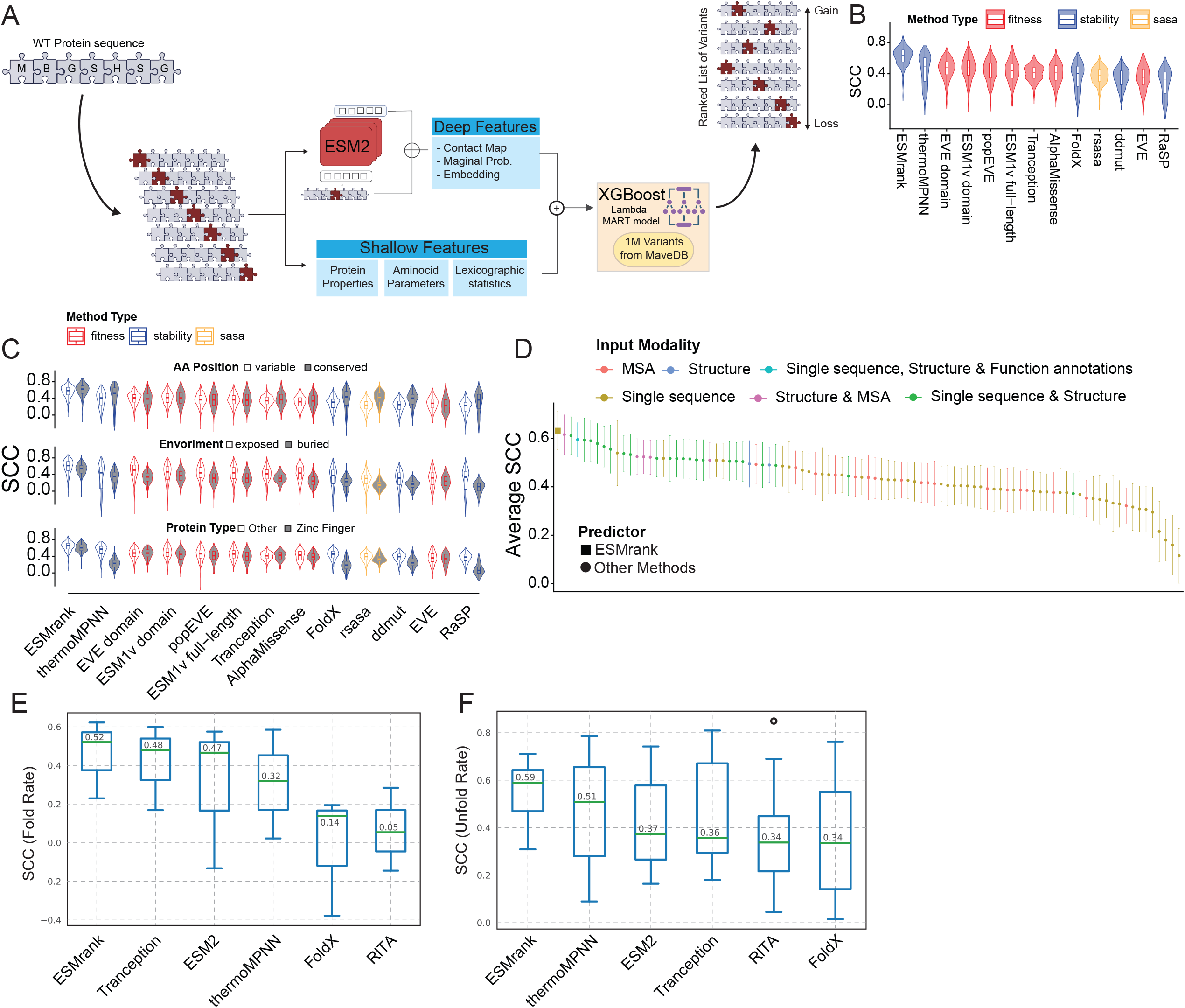
ESMRank outperforms state-of-the-art predictors across stability benchmarks. **(A)** Overview of the ESMRank architecture. Variants are encoded using deep representations derived from ESM models (embedding shifts, contact-map perturbations, masked marginal probabilities) together with curated biophysical and positional descriptors. These features are integrated within a LambdaMART learning-to-rank framework trained on ∼1 million soundness-normalized variants from MAVEdb to optimize the relative ordering of variants within each protein. **(B)** Benchmark on the Human Domainome dataset (∼500,000 mutations across ∼500 domains). Violin plots show the distribution of Spearman correlation coefficients (SCC) between predicted and experimental effects for multiple predictors. Domains overlapping with training proteins were excluded. ESMRank demonstrates superior agreement with experimental measurements across this large-scale benchmark. **(C)** Same as (B) but stratifying mutation structural and sequence contexts such as mutation occurring in variable (ConSurf score < 7) or conserved position (ConSurf score >=7) (upper plot) or exposed versus buried residues (middle panel) or protein family (bottom panel). **(D)** Performance on ProteinGym stability assays. Methods are grouped by input modality (sequence, structure, MSA, or combinations). ESMRank achieves the highest average ranking accuracy across datasets. Bars represent 95% confidence interval of the average Spearman Correlation Coefficient (n=14). **(E)** Evaluation on the VariBench folding benchmark. Spearman correlations are shown for prediction of folding rate constants, with ESMRank outperforming alternative approaches. (**F**) Same as (E) but for unfolding rate constants.

Across cross-validation, the learning-to-rank formulation consistently outperformed regression models trained on identical features, demonstrating that directly optimizing ordinal relationships improves discrimination of mutational impact across heterogeneous assay types (Fig. S4). Ablation analyses further showed that removing either deep or shallow feature groups resulted in a loss of ranking performance (Fig. S4), confirming that both sources of information contribute non-redundant signal. Feature-importance analysis on the full model revealed that the most informative predictors comprise a mixture of deep and shallow representations, with melting temperature and instability index among the strongest shallow features and masked marginal residue-probability shifts and embedding distances among the strongest deep features (Fig. S5).

Together, these results demonstrate that variant prioritization can be learned directly from protein sequence using a unified ranking framework that integrates local physicochemical perturbations with global, sequence-encoded structural context. This underscores the complementary value of protein language model representations and classical biophysical descriptors for modeling mutational effects.

### 3. ESMRank outperforms state-of-the-art predictors across stability benchmarks

To assess predictive performance and biological validity, we evaluated ESMRank on large-scale datasets measuring mutational effects on protein stability and abundance across diverse in vitro assays.

We first benchmarked ESMRank on the Human Domainome dataset (32), which quantifies the effects of 563,534 missense variants across 522 protein domains spanning all major structural classes, including metal-binding zinc-finger domains, using an abundance protein fragment complementation assay (aPCA), a cell-based readout that serves as a proxy for protein stability. Although the Domainome was not included in the MAVEdb training release, we ensured unbiased evaluation by excluding domains sharing >80% sequence identity with any training protein (Methods). ESMRank achieved a median Spearman correlation of 0.62, substantially outperforming ThermoMPNN (33) (ρ = 0.46) and other widely used stability and fitness predictors (Fig. 2B). Performance remained stable under increasingly stringent homology filtering (<50% and <25% identity; Fig. S6), demonstrating robust generalization beyond closely related proteins. Performance gains were consistent across conserved and variable positions, buried and surface-exposed residues, and zinc-finger domains (Fig. 2C), a class in which stability prediction is particularly challenging due to metal coordination requirements (32).

Importantly, these performance improvements reflected biologically meaningful predictions. ESMRank recapitulated the same structural organization observed in the integrated MAVEdb landscape (Fig. 1D–H): buried residues were less tolerant to substitution than exposed positions, nonsense mutations were uniformly deleterious, and hydrophobic-to-polar or charged substitutions were strongly destabilizing in protein cores (Fig. S7).

We next evaluated ESMRank on curated mutational scanning datasets from ProteinGym (13) in the zero-shot setting. After excluding proteins present in training and restricting evaluation to proteins sharing <80% sequence identity with any training protein, ESMRank was benchmarked against methods reported on the ProteinGym leaderboard. On stability-related assays, ESMRank achieved the highest mean Spearman correlation among all competing approaches (mean SCC = 0.63), surpassing sequence-based, structure-based, and hybrid predictors (Fig. 2D and Supplementary Table 02). To assess robustness across diverse contexts, we constructed six stratified evaluation subsets based on evolutionary conservation, intrinsic disorder, and residue exposure (Methods). Variants were evaluated separately in conserved versus variable regions, ordered versus intrinsically disordered regions, and buried versus surface-exposed positions. ESMRank ranked first in four of six strata, including variable and intrinsically disordered regions where many predictors show substantial performance degradation (Fig. S8). Across additional assay categories, including activity, expression, and organismal fitness, ESMRank remained competitive among sequence-only methods, while multimodal predictors leveraging structural models or multiple sequence alignments achieved higher performance in some settings (Supplementary Table 02). Notably, ESMRank’s stability performance and cross-context robustness were achieved using primary sequence alone.

Finally, to evaluate performance against independent kinetic measurements, we tested ESMRank on the VariBench (34) folding dataset (Methods). Across 1,273 mutations in 39 proteins, predictions correlated with both folding rates (median ρ = 0.52) and unfolding rates (median ρ = 0.59), outperforming ThermoMPNN (33), FoldX (35), and other predictors (Fig. 2E,F). These correlations recapitulate expected biophysical signatures of stabilizing and destabilizing substitutions, providing orthogonal kinetic validation.

Together, these results demonstrate that ESMRank delivers state-of-the-art performance on stability-focused benchmarks while maintaining robustness across structural and evolutionary contexts. By combining a learning-to-rank objective with sequence-derived structural representations, ESMRank captures fine-grained biophysical signals and enables accurate stability prediction from protein sequence alone.

### 4. ESMRank reveals mechanistic gradients of variant pathogenicity

The human proteome harbors a vast repertoire of missense variants whose clinical consequences arise through diverse molecular mechanisms, ranging from subtle modulation of activity to complete loss of function. Although impaired stability is a major contributor to pathogenic variation (32,36), its influence varies across structural environments and genetic contexts. Because ESMRank prioritizes mutations according to functional impact learned from heterogeneous experimental assays, we asked how its predictions align with clinical variant interpretation.

We first examined variants within the Human Domainome dataset, which includes 3,652 clinically annotated missense substitutions across 114 protein domains (Methods). Among these, 621 are classified as pathogenic or likely pathogenic and 322 as benign or likely benign, with the remainder annotated as variants of uncertain significance. Compared with experimental abundance measurements and established stability predictors, ESMRank produced a sharper separation between pathogenic and putative benign population variants from gnomAD (Fig. 3A), whereas ΔΔG-based approaches showed broader overlap between classes. Receiver-operating-characteristic analysis supported this trend: although AlphaMissense achieved the highest overall performance (AUC = 0.92), ESMRank was the strongest method among stability-oriented predictors (AUC = 0.78), substantially exceeding ThermoMPNN (AUC = 0.61) and FoldX (AUC = 0.63) (Fig. 3B). Importantly, ESMRank was not trained or fine-tuned on clinical labels and relies exclusively on sequence-derived features, demonstrating that clinically relevant constraint emerges from experimentally learned mutational ordering rather than explicit disease supervision.

**Figure 3.**
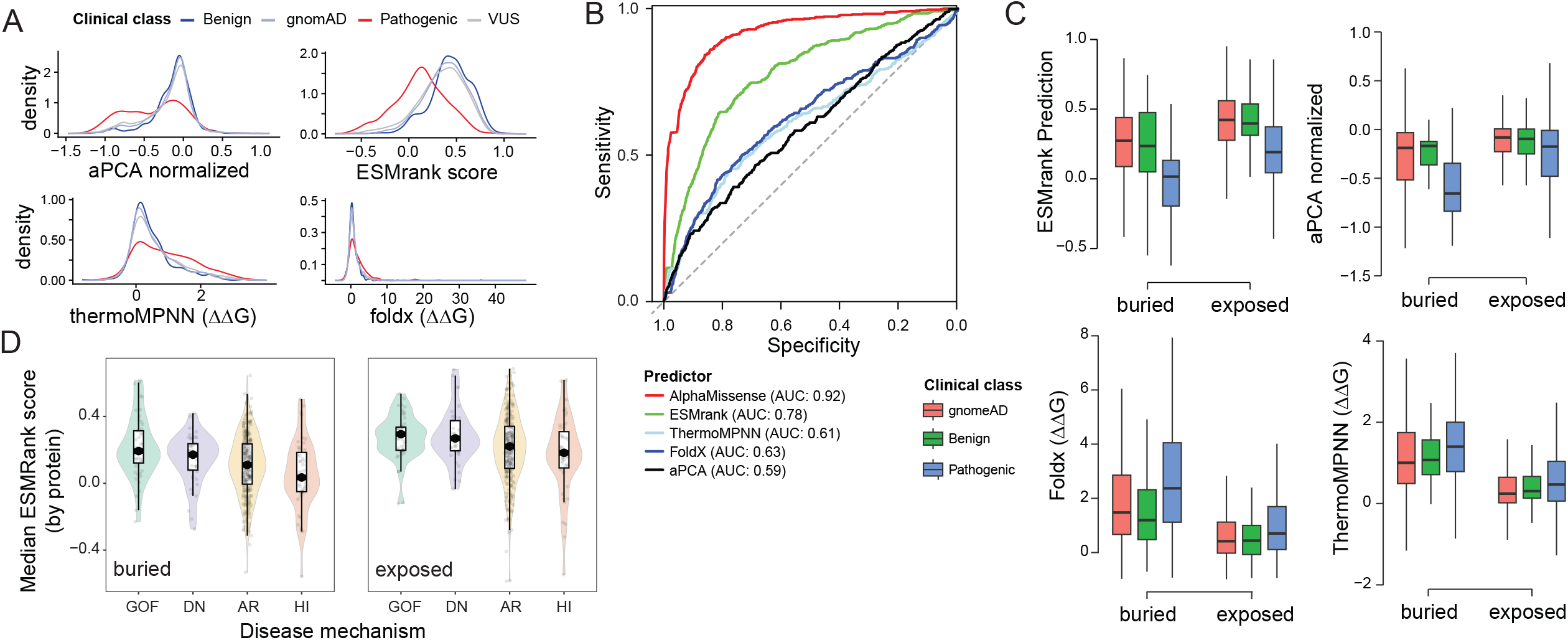
ESMRank captures clinically relevant gradients of variant pathogenicity. (**A**) Distributions of experimental abundance scores (aPCA) and computational predictions for clinically annotated missense variants from the Human Domainome dataset, stratified as pathogenic, benign, putatively benign population variants from gnomAD, or variants of uncertain significance (VUS). ESMRank shows improved separation between pathogenic and benign classes compared with ΔΔG-based predictors. (**B**) Receiver operating characteristic curves for discrimination of pathogenic versus benign variants. AlphaMissense achieves the highest overall performance (AUC = 0.92), whereas ESMRank is the strongest performer among stability-oriented approaches (AUC = 0.78), exceeding ThermoMPNN, FoldX, and experimental abundance measurements. (**C**) Variant effects stratified by solvent exposure. Pathogenic variants are more deleterious than benign variants at both buried and exposed positions. Whereas ΔΔG-based predictors show reduced separation at exposed residues, ESMRank maintains discriminatory power across environments. (**D**) Protein-level aggregation of ESMRank scores reveals a mechanistic gradient across genes associated with different modes of pathogenic action. Median mutational tolerance is highest for gain-of-function (GOF) genes, followed by dominant-negative (DN), autosomal recessive (AR), and haploinsufficient (HI) genes. Separation is more pronounced for buried than exposed residues.

We next investigated how structural environment modulates this relationship. Stratifying ClinVar and gnomAD (37) variants by solvent exposure showed that pathogenic substitutions were, on average, more deleterious than benign ones at both buried and exposed sites. However, whereas the discriminatory power of ΔΔG-based predictors decreased substantially at exposed positions, ESMRank maintained separation across both environments (Fig. 3C). Similar trends were observed in an expanded analysis spanning ∼12,000 proteins and approximately 95,000 ClinVar missense variants, alongside ∼8,000 common gnomAD variants used as benign proxies (Fig. S9). These results indicate that, alongside classical stability effects dominant in protein cores, the information captured by ESMRank remains informative in regions where explicit thermodynamic proxies become less predictive.

Finally, we tested whether genes associated with distinct pathogenic mechanisms exhibit systematic differences in their global mutational tolerance. We analyzed 8,081 pathogenic ClinVar missense variants curated by Ziang Li et al. (38), spanning 836 genes classified into four mutually exclusive mechanistic categories: gain-of-function (GOF), dominant-negative (DN), autosomal recessive (AR), and haploinsufficiency (HI). Because pathogenic mechanism is fundamentally a gene-level property, we aggregated ESMRank scores per protein, stratified by structural environment. This revealed a robust and graded constraint landscape aligned with expected levels of residual molecular activity. Genes associated with GOF disease exhibited the highest overall tolerance, followed by DN and AR genes, whereas HI genes were the most constrained (Fig. 3D). As anticipated, substitutions at exposed positions were generally more permissive than those in the core, with mechanistic stratification most pronounced among buried residues. Notably, this organization emerges despite the absence of inheritance or phenotypic information during training, suggesting that constraints relevant to disease mechanisms are implicitly encoded in experimentally derived mutational patterns.

Together, these results indicate that ESMRank captures a continuous, mechanism-aware spectrum of pathogenicity in which stability-related effects represent a prominent, but not exclusive, axis of disease causation. Rather than serving as a generic pathogenicity predictor, these observations suggest that ESMRank captures stability-mediated components of pathogenicity in a context-dependent manner, motivating focused analysis of diseases in which protein instability is a dominant and therapeutically relevant mechanism.

### 5. ESMRank links CFTR structural constraint to functional impairment and therapeutic responsiveness

Cystic fibrosis (CF) arises from missense variants in the cystic fibrosis transmembrane conductance regulator (CFTR) that disrupt folding, trafficking, and channel gating (39). CFTR therefore provides a paradigmatic system in which protein stability directly influences both disease severity and therapeutic responsiveness. We used CFTR as a case study to test whether ESMRank captures stability-mediated constraints that underlie pathogenicity and pharmacological rescue.

Mapping per-residue ESMRank scores onto the CFTR structure (PDBID 5UAK) (40) revealed marked intolerance to substitution within transmembrane helices and interdomain interfaces (Fig. 4A,B). Along the primary sequence, ESMRank exhibits pronounced positional heterogeneity, with lowest scores localizing to TMD1, TMD2, and the nucleotide-binding domains, regions known to be structurally constrained and sensitive to destabilizing mutations. Comparison with evolutionary conservation (Fig. 4B) shows that several of these regions display only moderate conservation, indicating that ESMRank captures biophysical constraints related to membrane insertion, helix packing, and domain coupling that are not fully reflected by sequence conservation alone.

**Figure 4.**
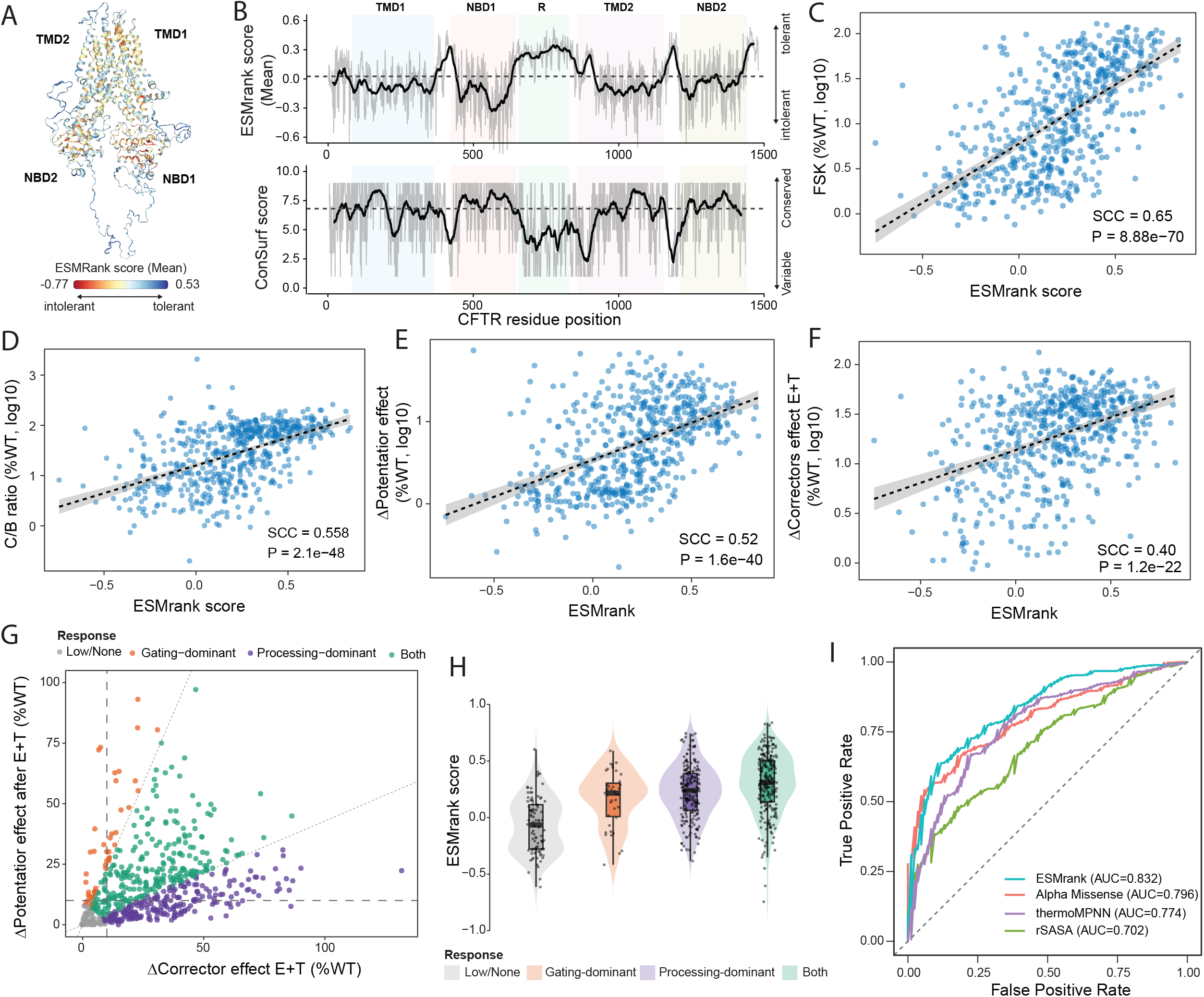
ESMRank links CFTR structural constraint, functional output, and therapeutic responsiveness. (**A)** Structural mapping of per-residue ESMRank scores onto the CFTR cryo-EM structure. Regions predicted to be intolerant to substitution (red spots) localize predominantly to transmembrane helices and interdomain interfaces, including TMD1, TMD2, NBD1, and NBD2. (**B**) Positional distribution of ESMRank (top) and ConSurf evolutionary conservation scores (bottom) along the CFTR sequence. ESMRank reveals pronounced troughs within structurally constrained domains (TMD1, NBD1, R, TMD2, NBD2), including regions with only moderate evolutionary conservation, indicating sensitivity to biophysical constraints not fully captured by sequence conservation alone. (**C)** Correlation between ESMRank score and basal CFTR channel function measured as transepithelial conductance (FSK, %WT, log□ □ scale) across 585 missense variants (Spearman ρ = 0.65, P = 8.88×10□ □ □). (**D**) Correlation between ESMRank score and maturation efficiency quantified by the ratio of fully glycosylated to immature CFTR (C/B ratio, %WT, log□ □ scale) (Spearman ρ = 0.558, P = 2.1×10□ □ □). (**E)** Association between ESMRank score and potentiator (ivacaftor, IVA) response (Δpotentiator effect, %WT, log□ □ scale) (Spearman ρ = 0.52, P = 1.6×10□ □ □). (**F**) Association between ESMRank score and dual corrector (elexacaftor + tezacaftor; ELX+TEZ) response (Δcorrector effect, %WT, log□ □ scale) (Spearman ρ = 0.40, P = 1.2×10□^22^). Dashed lines in C–F indicate linear regression fits with 95% confidence intervals. (**G**) Mechanistic classification of variants based on relative contributions of corrector and potentiator effects. Variants are categorized as Low/None, Gating-dominant, Processing-dominant, or Both (responsive to both modulators). (**H**) Distribution of ESMRank scores across response classes. Variants responsive to both modulators exhibit the highest median ESMRank scores, whereas low/non-responsive variants show the lowest scores, indicating a graded relationship between predicted sequence constraint and therapeutic responsiveness. (**I**) Receiver operating characteristic (ROC) curves for classification of therapeutically responsive variants. ESMRank (AUC = 0.832) outperforms AlphaMissense (AUC = 0.796), thermoMPNN (AUC = 0.774), and a solvent accessibility baseline (rSASA; AUC = 0.702).

We next evaluated ESMRank against clinically annotated CFTR missense variants from the CFTR2 database (41). Restricting the analysis to CF-causing and non–CF-causing variants, ESMRank achieved robust discrimination (ROC AUC = 0.92), approaching the performance of AlphaMissense (ROC AUC = 0.97) while outperforming ΔΔG-based predictors such as ThermoMPNN (ROC AUC = 0.7) and FoldX (ROC AUC = 0.7) (Fig. S10). Notably, ESMRank was not trained on CFTR-specific data or clinical labels, indicating that pathogenic signal emerges from general sequence-derived constraints rather than disease-specific supervision.

To assess whether ESMRank reflects mechanistic determinants of CFTR biogenesis, we analyzed 585 missense variants (42) from a comprehensive theratype screen measuring basal channel function, maturation efficiency, and pharmacological response to the correctors elexacaftor (ELX) and tezacaftor (TEZ), and the potentiator ivacaftor (IVA). ESMRank correlated strongly with basal transepithelial conductance (Spearman ρ = 0.65, P = 8.88×10□ □ □; Fig. 4C) and with maturation efficiency measured by C/B ratio (ρ = 0.558, P = 2.1×10□ □ □; Fig. 4D), supporting a direct relationship between predicted sequence constraint and folding efficiency.

ESMRank further correlated with pharmacological responsiveness: IVA potentiator effect (ρ = 0.52, P = 1.6×10□ □ □; Fig. 4E) and dual corrector (ELX+TEZ) response (ρ = 0.40, P = 1.2×10□^22^; Fig. 4F) both increased with higher ESMRank scores. Consistent associations were independently observed in the dataset reported by McKee et al. (43), which evaluated CFTR abundance and corrector responses across 129 missense variants. ESMRank correlated with baseline CFTR abundance (ρ = 0.49; P = 2.63×10^-7^; Fig. S11A), ELX response (ρ = 0.52; P = 2.5×10^-8^; Fig. S11B), TEZ response (ρ = 0.51; P = 5.7×10^-8^; Fig. S11C), and their combined effect (ρ = 0.54; P = 6.21×10^-9^; Fig. S11D). These concordant trends indicate that the association between predicted sequence constraint and pharmacological responsiveness is reproducible across experimental systems. Collectively, the upward relationships observed across panels support a model in which variants predicted to be less destabilizing retain sufficient structural integrity to enable both correction of folding defects and potentiation of channel activity.

To further dissect therapeutic mechanisms, we classified variants into four response categories based on the relative contributions of corrector and potentiator effects (Fig. 4G). Gating-dominant variants exhibited low Δcorrector but high Δpotentiator responses, whereas processing-dominant variants showed the reciprocal pattern; mixed variants occupied the upper-right quadrant, reflecting responsiveness to both modulators. These mechanistic classes displayed graded shifts in ESMRank distributions (Fig. 4H and Methods), with low or non-responsive variants showing the lowest median scores, gating- and processing-dominant variants occupying intermediate ranges, and mixed variants exhibiting the highest scores. This stratification indicates that ESMRank captures a stability-centered continuum that modulates both folding competence and pharmacological tractability. Consistent with this, ROC analysis showed that ESMRank achieved the highest discrimination (AUC = 0.83), exceeding AlphaMissense (AUC = 0.80) and ThermoMPNN (AUC = 0.77) (Fig. 4I). Moreover, stratification by mechanistic response class revealed pronounced differences in subclass performance, with AlphaMissense showing limited sensitivity for gating-dominant variants (AUC = 0.53), whereas ESMRank always retained discriminatory power across all mechanistic subclasses (mean ESMRank AUC = 0.82 vs mean AlphaMissense AUC = 0.73) (Fig. S12).

Together, these results indicate that ESMRank captures a stability-centered axis of CFTR variant effects that links pathogenicity, folding efficiency, and therapeutic tractability. Importantly, this signal emerges without CFTR-specific training or pharmacological annotations, demonstrating that clinically relevant structure–function relationships can be inferred directly from general sequence-derived constraints. More broadly, these findings suggest that stability-informed sequence models may provide a general framework for anticipating both disease mechanisms and pharmacological responsiveness across genetically heterogeneous disorders.

## DISCUSSION

Multiplexed assays of variant effect (MAVEs) have generated extensive maps of mutational tolerance across diverse proteins and experimental systems. However, differences in assay design, scale, and biological context have limited systematic integration across studies. Here we show that partial redundancy among independent experiments can be leveraged to recover reproducible ordinal structure in mutational landscapes. By quantifying cross-experiment rank consistency, our overlap-aware framework derives a consensus measure of variant tolerance that reduces assay-specific scaling differences while preserving within-protein ordering relationships. Applied to more than two million missense variants, this strategy reveals coherent structural and biochemical gradients consistent with known determinants of protein stability and organization.

An important feature of the recovered landscape is its enrichment for stability-associated constraint. This likely reflects not only the composition of current MAVE datasets, which include numerous abundance- and folding-proxy assays, but also a more general property of protein biology. Variants that substantially impair folding reduce steady-state protein abundance and thereby limit participation in downstream biochemical processes. As a consequence, severe folding defects can manifest as deleterious effects across diverse assay modalities, including measurements of activity, interaction, and organismal fitness (16). Because the overlap-aware integration strategy emphasizes rank consistency across experiments, variants that reproducibly impair protein folding across multiple contexts are preferentially reinforced. In this sense, the dominant axis recovered here may reflect a common biophysical bottleneck that persists across heterogeneous experimental systems rather than a bias intrinsic to any single assay type. At the same time, this stability-influenced dimension does not necessarily exhaust the spectrum of mutational constraint encoded across functional assays. Context-specific effects such as regulatory modulation, interaction rewiring, or environment-dependent activity changes may contribute additional, partially orthogonal dimensions that are less consistently captured across current datasets. As the diversity of MAVE modalities expands, it will be important to determine whether overlap-aware integration reveals additional reproducible constraint dimensions beyond the stability-centered signal described here.

Our findings indicate that transferable mutational signal resides primarily in relative variant ordering rather than absolute effect magnitude. We show that jointly aligning model training and representation with evolutionary and biophysical constraints improves generalizable prediction of variant effect. Optimizing ranking performance through a learning-to-rank formulation, while integrating protein language model embeddings with complementary physicochemical descriptors, enabled the development of ESMRank, a sequence-based predictor that performs strongly on large-scale stability and abundance benchmarks under strict protein-level partitioning. Protein language models encode rich evolutionary and contextual information, including implicit structural priors, but do not explicitly parameterize thermodynamic determinants such as hydrophobicity, packing density, or aggregation propensity. Conversely, classical physicochemical features capture interpretable stability-related axes that are only partially reflected in learned embeddings. Their integration increases robustness under heterogeneous experimental supervision, indicating that evolutionary representations and explicit biophysical priors provide complementary inductive biases for modeling mutational constraint. Although ESMRank was not trained on clinical annotations, the reconstructed constraint landscape is enriched for pathogenic variants and stratifies genes by established disease mechanisms, demonstrating that reproducible, stability-linked components of mutational constraint are encoded in sequence and recoverable through representation learning.

The CFTR case study illustrates how sequence-derived mutational tolerance can align with experimentally measured folding efficiency, channel function, and pharmacological responsiveness. Although this analysis focuses on a protein in which stability plays a central pathogenic role, it highlights the potential for integrated functional landscapes to inform mechanistic interpretation in genetically heterogeneous disorders. Whether similar relationships extend to proteins dominated by non-stability-mediated mechanisms warrants further investigation.

Several additional considerations merit attention. First, while protein-level partitioning and sequence identity filtering reduce information leakage, generalization to remote structural classes will depend on continued diversification of experimental data. Second, the current framework models single-amino-acid substitutions independently and does not capture higher-order epistatic effects. Third, sequence-only modeling does not explicitly incorporate dynamic conformational changes or ligand-dependent states, which may influence variant impact in specific contexts.

Despite these limitations, our results demonstrate that heterogeneous MAVE datasets contain reproducible ordinal structure that can be extracted through overlap-aware integration and transferred through sequence-based modeling. Rather than replacing clinically supervised predictors, this approach provides a complementary, experimentally grounded view of mutational constraint that emphasizes stability-associated components of variant impact. As functional genomics datasets continue to expand, principled integration strategies may help bridge fragmented experimental measurements with scalable, mechanistically informed variant prioritization.

