## Supplementary Figure for "ESMRank reveals a transferable axis of protein mutational constraint from overlapping variant effect assays"

**AFFILIATIONS**
[1]Telethon Institute of Genetics and Medicine, Naples, Italy
[2]University of Naples Federico II, Department of Electrical Engineering and Information Technology, Naples, Italy
[3]Scuola Superiore Meridionale, Genomics and Experimental Medicine Program, Naples, Italy

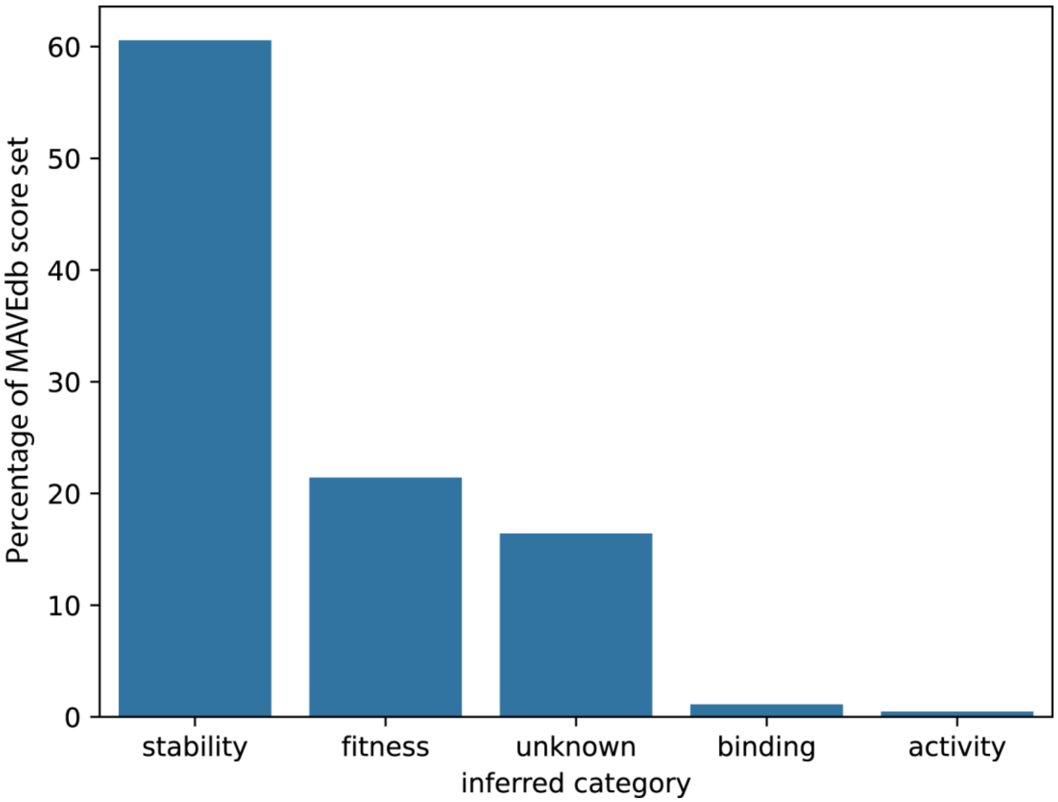


**Figure S1 | Functional composition of MaveDB score sets used for overlap-aware fusion.** Distribution of MaveDB experiments by inferred functional category following preprocessing and filtering. Categories were assigned through a metadata-based survey of assay descriptions using keyword matching. The majority of retained score sets correspond to protein stability assays (~60%), followed by fitness-related assays (~21%) and experiments with ambiguous or composite functional readouts (~17%). Binding (~1%) and activity (<1%) assays represent a minor fraction of the dataset. Percentages are calculated relative to the total number of MaveDB score sets included in the overlap-aware integration procedure used in this study.

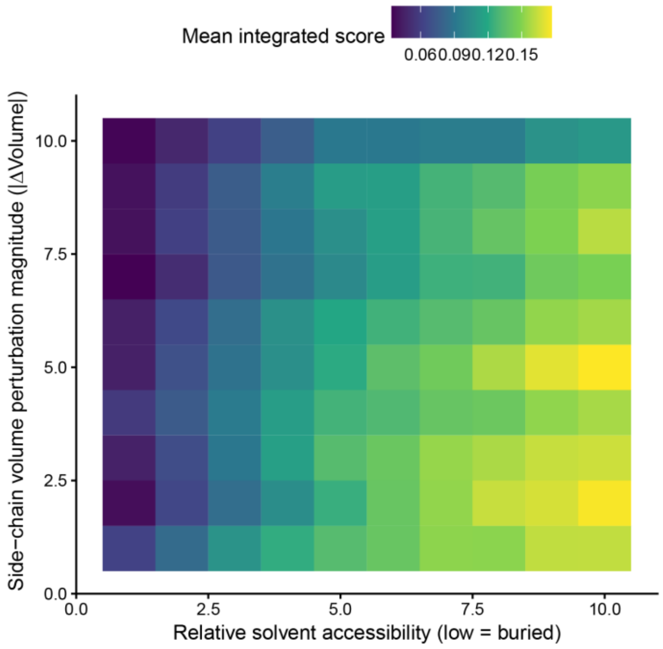


**Figure S2 | Quantitative interaction between solvent accessibility and side-chain volume perturbation in the integrated mutational landscape.** Heatmap showing the mean integrated (soundness-normalized) variant score across joint bins of relative solvent accessibility (rSASA) and side-chain volume perturbation magnitude (|ΔVolume|). rSASA was computed from AlphaFold structural models using per-atom solvent-accessible surface area calculated with FreeSASA and normalized by residue-specific maximal solvent accessibility values (SASA_max). Per-residue rSASA was obtained by summing atomic contributions and dividing by SASA_max for the corresponding amino acid. Only residues with mean per-residue pLDDT ≥ 70 were included. Side-chain volume perturbation was defined as the absolute difference between wild-type and mutant residue side-chain volumes (|ΔVolume|), using established amino acid volume parameters. Variants were stratified independently into deciles of rSASA and |ΔVolume| using empirical quantiles across all substitutions. For each of the resulting 10 × 10 bins, the mean integrated score was computed. The analysis included 650,505 single–amino-acid substitutions across 533 proteins. Mean mutational tolerance increases monotonically with solvent exposure (left to right) and decreases with increasing perturbation magnitude (bottom to top). The lowest tolerance values occur at the intersection of low rSASA (buried residues) and high |ΔVolume| (large packing perturbations), consistent with steric destabilization effects in the protein core. To formally test this interaction, a linear mixed-effects model with protein-level random intercepts was fitted, with exposure class defined as buried (rSASA < 0.25) or surface (rSASA ≥ 0.25). The interaction term was highly significant ($t = -16.05$), indicating that the impact of side-chain volume perturbation on mutational tolerance is significantly stronger at buried positions than at surface-exposed residues. These results suggest that the integrated mutational axis encodes graded structural constraints consistent with packing-mediated stability effects rather than simple categorical core–surface differences.

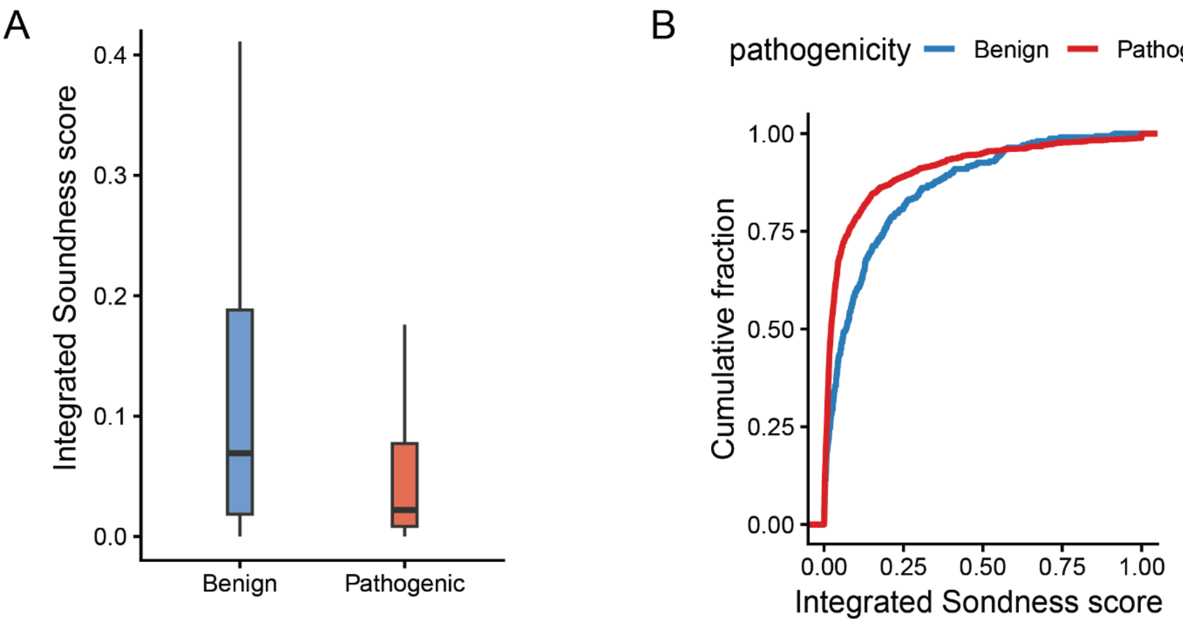


**Figure S3 | Integrated mutational soundness distinguishes ClinVar pathogenic and benign variants.** **(A)** Distribution of integrated (soundness-normalized) scores for ClinVar missense variants annotated as benign/likely benign (n = 310) or pathogenic/likely pathogenic (n = 1,069) after intersection with the MaveDB-integrated dataset at the protein level. Pathogenic variants are significantly enriched at lower (more deleterious) integrated scores compared to benign variants (two-sided Wilcoxon rank-sum test, $P = 2.0 \times 10^{-12}$). Boxes represent the interquartile range (IQR), center lines indicate medians, and whiskers extend to 1.5× IQR. **(B)** Empirical cumulative distribution functions (ECDFs) of integrated scores for benign (blue) and pathogenic (red) variants. Pathogenic variants exhibit a pronounced leftward shift toward lower soundness values, consistent with enrichment of loss-of-function effects in human genetic disease. Together, these results demonstrate that the unified mutational landscape captures clinically relevant constraint patterns that generalize beyond experimental datasets to independent human pathogenic variation.

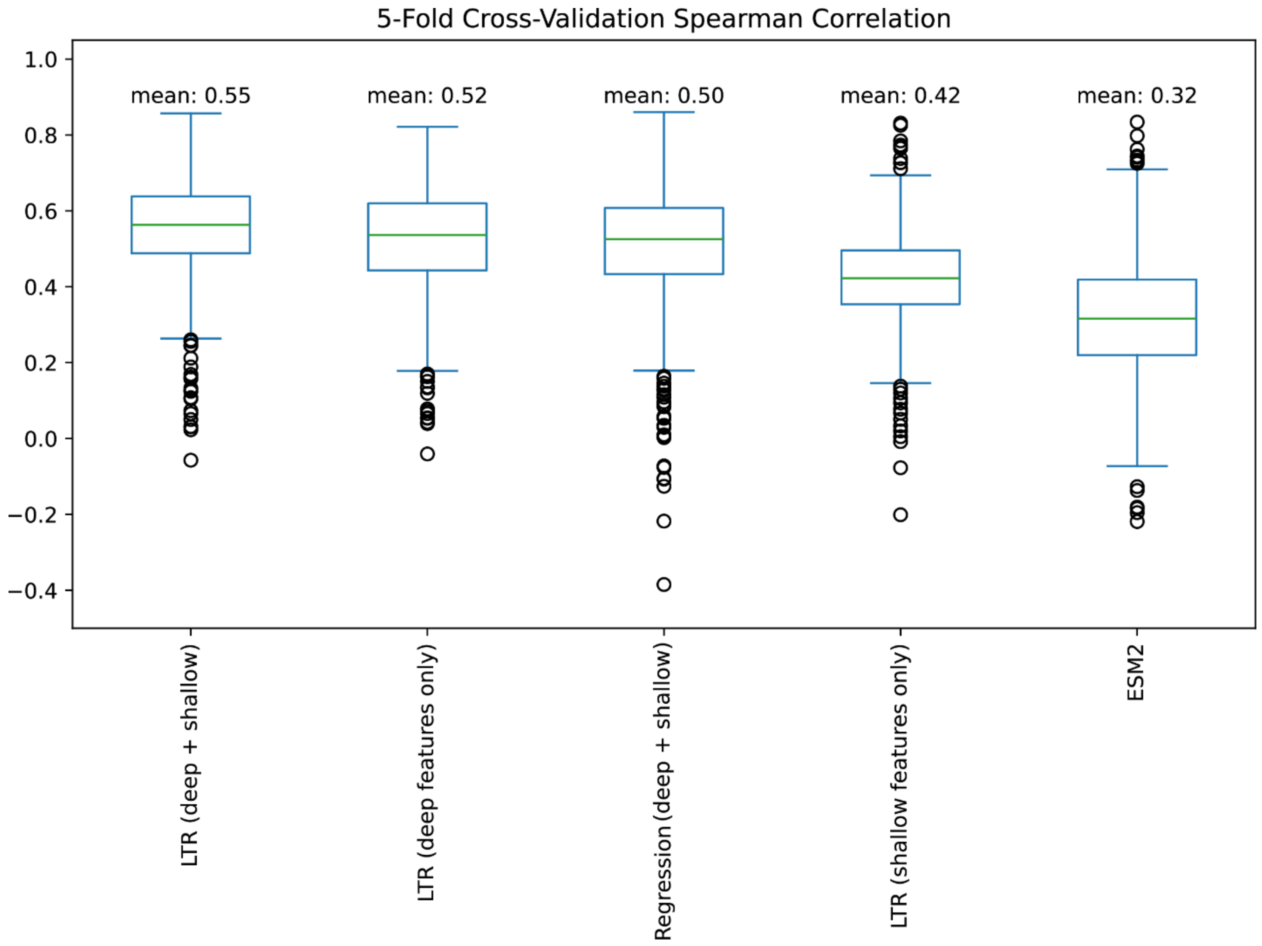


**Figure S4 | Cross-validation performance and feature ablation analyses.** Distribution of per-assay Spearman correlations obtained from 5-fold cross-validation under different model formulations and feature configurations. Shown are learning-to-rank (LTR) models trained on combined deep and shallow features, deep features only, or shallow features only; a regression model trained on identical combined features; and ESM2 as a baseline. Mean correlation values across folds are indicated above each box. The learning-to-rank formulation using both deep and shallow features achieved the highest average performance (mean $\rho = 0.55$), outperforming regression trained on the same feature set (mean $\rho = 0.50$) and ESM2 (mean $\rho = 0.32$). Ablation of either deep or shallow feature groups reduced ranking performance (deep only: mean $\rho = 0.52$; shallow only: mean $\rho = 0.42$), indicating that both feature classes contribute complementary, non-redundant signal. Boxes represent interquartile ranges, center lines indicate medians, whiskers extend to 1.5× IQR.

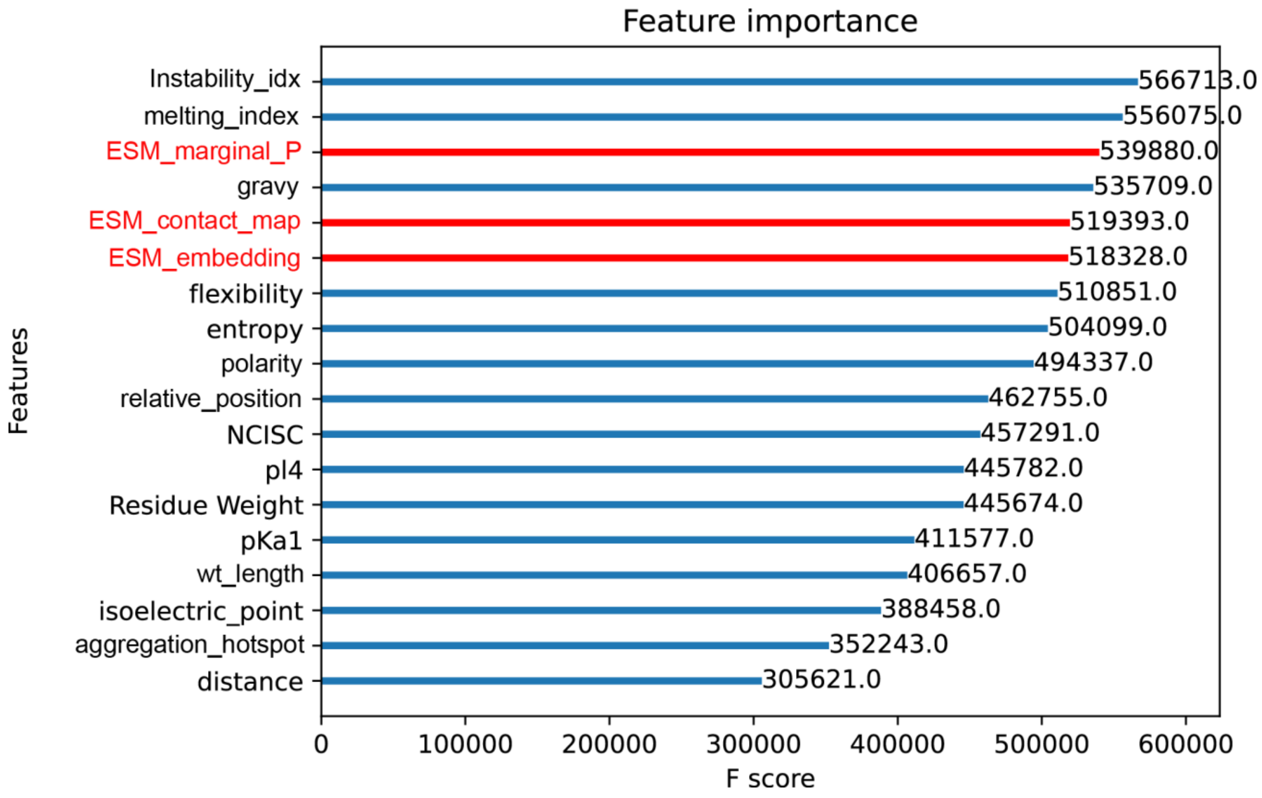


**Figure S5 | ESMRank full model feature importance.** Relative feature importance scores from the final LambdaMART model underlying ESMRank. Importance values correspond to cumulative split gain ("F score") across boosted decision trees and reflect each feature's contribution to ranking performance during training. Features are ordered by decreasing importance. Deep features derived from ESM2 representations (masked marginal probability, contact-map–based structural metric, and embedding distance between wild-type and mutant sequences; highlighted in red) rank among the top contributors, alongside key physicochemical descriptors such as instability index, predicted melting index, and hydropathy (GRAVY). Both deep and shallow features contribute substantially, supporting a complementary integration of learned sequence representations and classical biophysical properties in modeling mutational constraint.

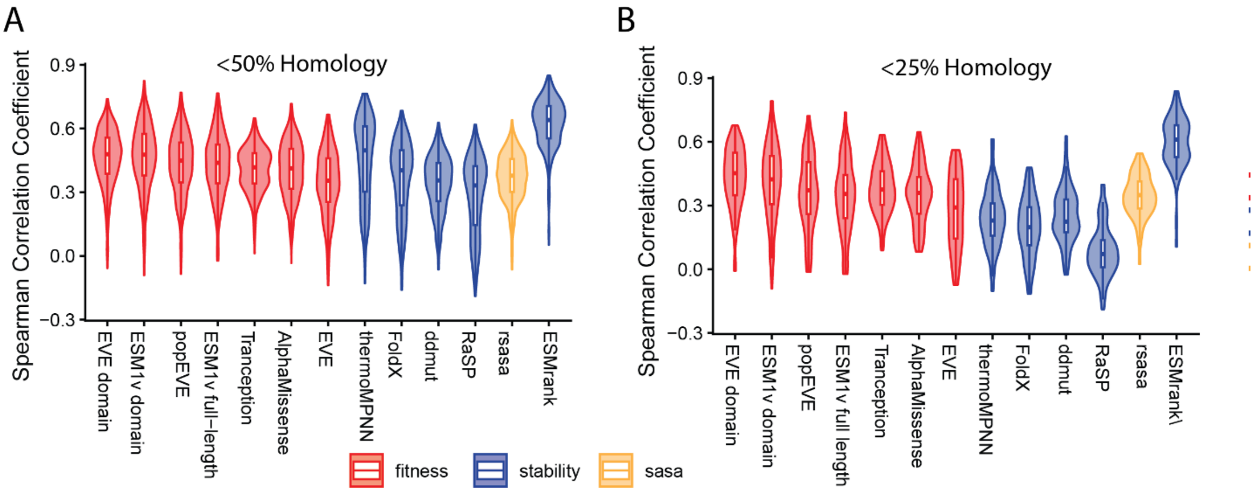


**Figure S6 | Performance under stringent homology filtering on the Human Domainome benchmark.** **(A)** Distribution of per-domain Spearman correlations for variants in domains sharing <50% sequence identity with any protein in the training set. Predictors are grouped by input modality: sequence-based fitness models (red), structure-based stability models (blue), and solvent accessibility baseline (orange). ESMRank maintains the highest median performance across assay types under this homology threshold. **(B)** Same analysis restricted to domains sharing <25% sequence identity with any training protein. Despite the increased evolutionary distance, ESMRank retains strong predictive performance, whereas most alternative predictors exhibit more pronounced performance degradation. Across both homology thresholds, ESMRank consistently outperforms ThermoMPNN and other widely used stability and fitness predictors, demonstrating robust generalization beyond closely related proteins and confirming that performance is not driven by residual sequence similarity to the training set.

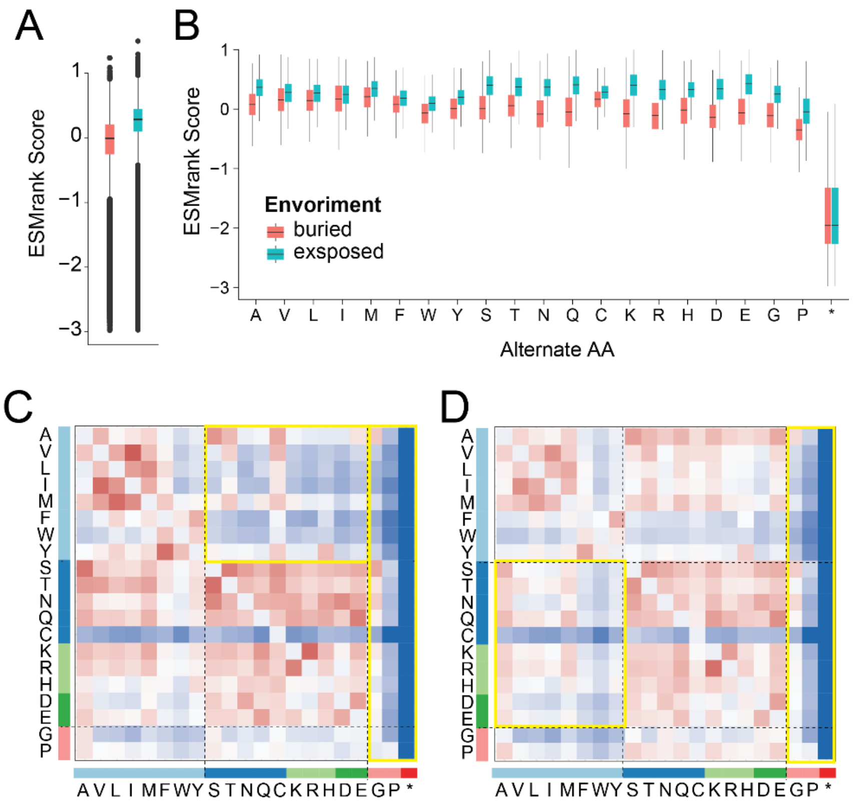


**Figure S7 | ESMRank recapitulates structural and physicochemical constraints observed in the integrated mutational landscape. (A)** Global distribution of ESMRank scores across all single–amino-acid substitutions in the Human Domainome dataset. Scores are oriented such that lower values indicate greater predicted deleteriousness. The distribution exhibits a pronounced deleterious tail, reflecting strong penalization of highly disruptive substitutions. **(B)** Distribution of ESMRank scores stratified by structural environment (buried vs exposed) and alternate amino acid identity. Substitutions at buried positions (red) are consistently more deleterious than those at exposed sites (teal) across most amino acid changes. Nonsense mutations (*) display uniformly low scores, indicating strong predicted functional impairment independent of structural context. **(C)** Mean substitution matrix of ESMRank scores for buried residues. Rows correspond to wild-type amino acids and columns to alternate residues. Hydrophobic-to-polar or charged substitutions in the protein core are strongly penalized, whereas conservative substitutions among hydrophobic residues show comparatively higher tolerance. The highlighted region emphasizes destabilizing substitutions that introduce polar or charged side chains into hydrophobic environments. **(D)** Mean substitution matrix for exposed residues. In contrast to buried positions, substitutions at solvent-exposed sites exhibit more moderate penalties and greater tolerance across physicochemical classes. The highlighted region illustrates the reduced constraint on polar or charged substitutions at surface positions. Together, these analyses demonstrate that ESMRank reproduces key structural and biochemical principles observed in the integrated MAVEdb landscape (Fig. 1D–H), including core–surface asymmetry, uniform intolerance of nonsense mutations, and strong destabilization of hydrophobic core packing by polar or charged substitutions.

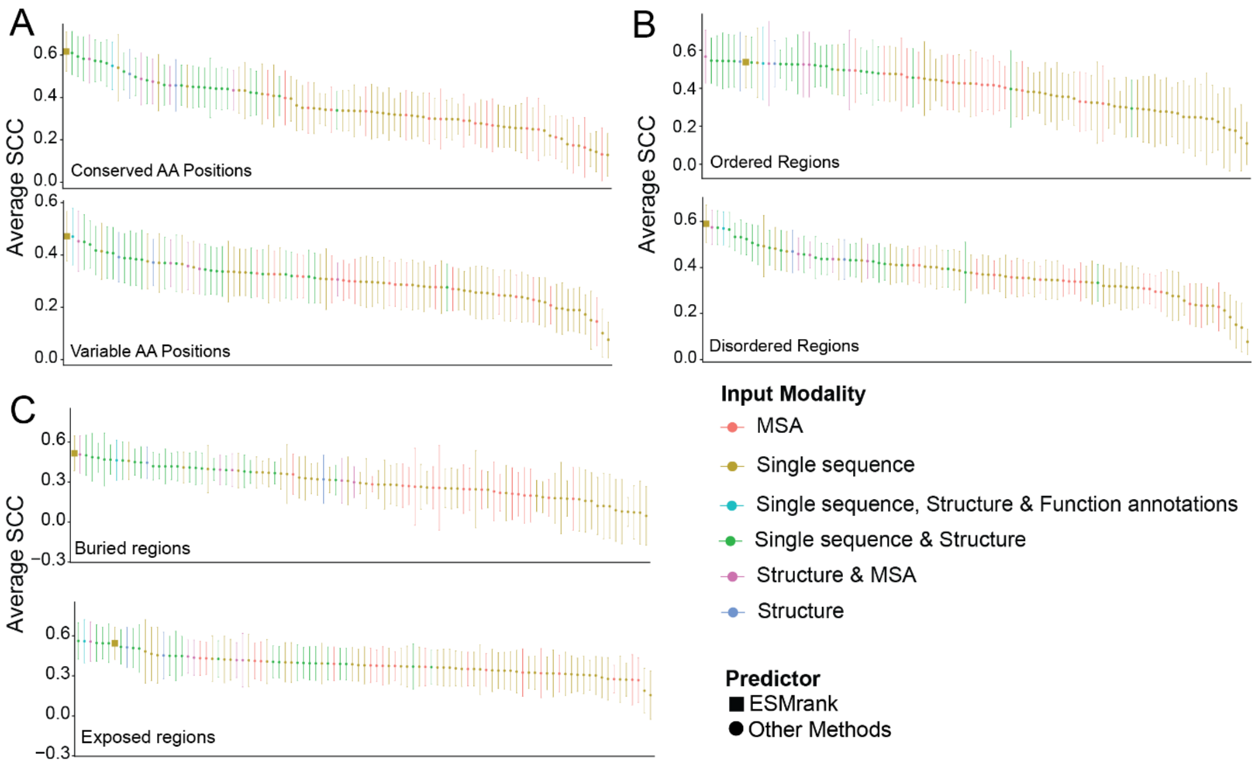


**Figure S8 | Stratified ProteinGym benchmarking across evolutionary and structural contexts.** Average per-assay Spearman correlation coefficients (SCC) for predictors evaluated on ProteinGym stability-related assays under stratified variant subsets. Error bars represent 95% confidence intervals of the mean Spearman correlation across assays. Colors denote input modality (MSA-based, single-sequence, structure-based, or hybrid methods), and square markers indicate ESMRank. **(A)** Performance in conserved versus variable amino acid positions, defined using ConSurf conservation scores. ESMRank ranks first in variable positions and remains competitive in conserved positions, indicating robust generalization beyond strongly constrained residues. **(B)** Performance in ordered versus intrinsically disordered regions, defined using IUPred2 predictions. ESMRank maintains leading performance in disordered regions, where many structure-dependent predictors exhibit reduced accuracy. **(C)** Performance in buried versus surface-exposed positions, defined using relative solvent accessibility computed from AlphaFold structural models. ESMRank performs strongly in both contexts and ranks first in buried residues, consistent with its ability to capture stability-related constraints. Across the six stratified contexts, ESMRank ranks first in four and remains competitive in the remaining two, demonstrating cross-context robustness despite relying solely on primary sequence input.

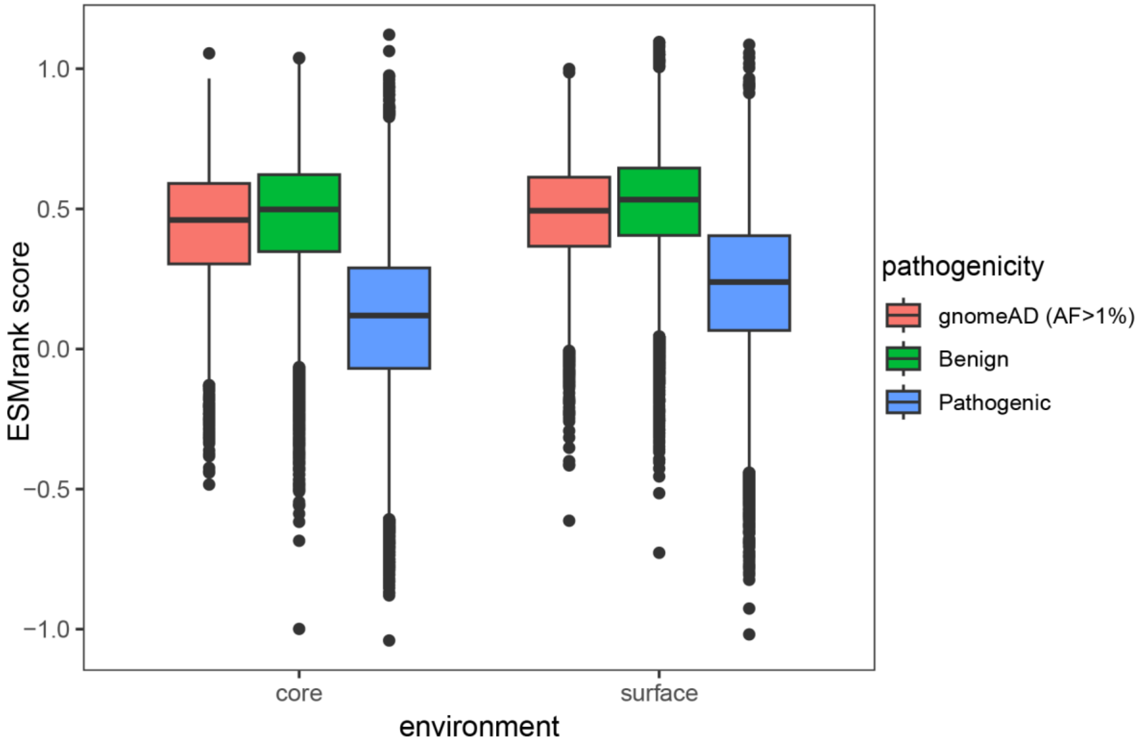


**Figure S9 | Structural context modulates pathogenicity discrimination by ESMRank.** Distribution of ESMRank scores for ClinVar pathogenic variants, ClinVar benign variants, and common gnomAD variants (allele frequency >1%) stratified by structural environment (core vs surface), defined using relative solvent accessibility computed from AlphaFold models. Scores are oriented such that lower values indicate greater predicted deleteriousness. In both buried (core) and exposed (surface) positions, pathogenic variants exhibit significantly lower ESMRank scores compared to benign and common population variants. Notably, separation between pathogenic and benign distributions is preserved at surface-exposed sites, where traditional ΔΔG-based stability predictors typically show reduced discriminatory power. This expanded analysis spans approximately 12,000 proteins and ~95,000 ClinVar missense variants, together with ~8,000 common gnomAD missense variants used as benign proxies, demonstrating that ESMRank captures biologically meaningful constraint signals beyond classical core stability effects.

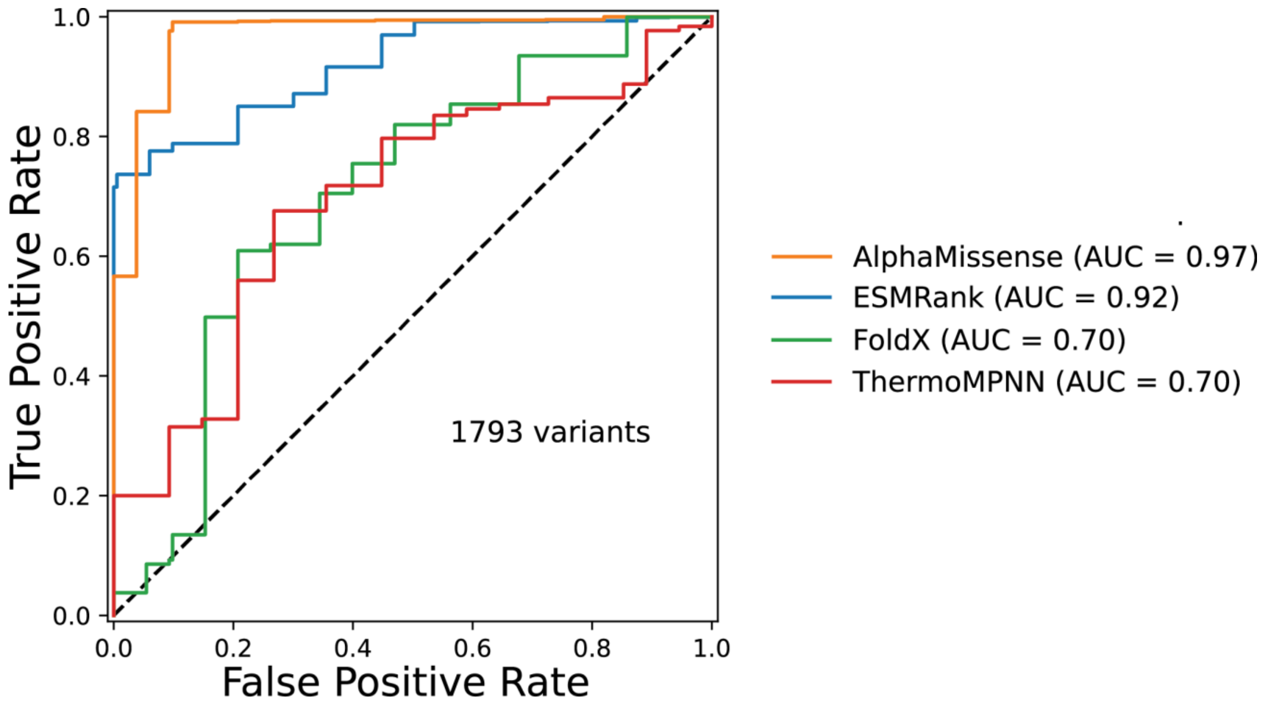


**Figure S10 | Discrimination of CFTR pathogenic variants.** Receiver operating characteristic (ROC) curves for classification of CF-causing versus non–CF-causing CFTR missense variants from the CFTR2 database (n = 1,793 variants). Performance is reported as area under the ROC curve (AUC). AlphaMissense achieved the highest discrimination (AUC = 0.97), followed by ESMRank (AUC = 0.92). ΔΔG-based stability predictors showed substantially lower performance (ThermoMPNN AUC = 0.70; FoldX AUC = 0.70). The dashed diagonal line indicates random classification. Importantly, ESMRank was neither trained on CFTR-specific data nor developed for pathogenicity prediction, indicating that clinically relevant pathogenic signal emerges from general sequence-derived constraints rather than disease-specific supervision.

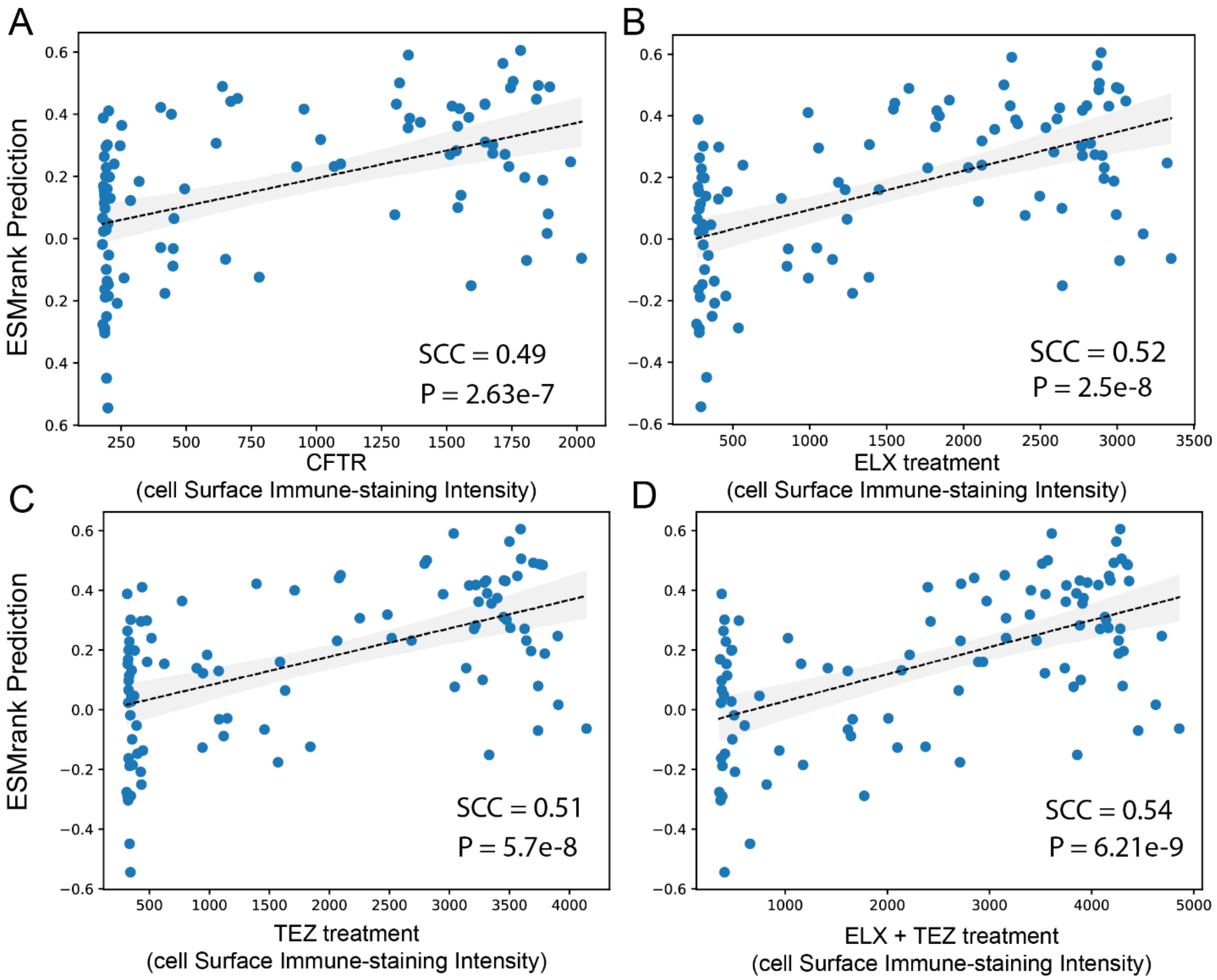


**Figure S11 | ESMRank correlates with CFTR abundance and corrector responsiveness in an independent mutational dataset.** Scatter plots showing the relationship between ESMRank predictions and experimental measurements of CFTR cell-surface abundance and pharmacological rescue across 129 missense variants (McKee et al.). Dashed lines indicate linear regression fits; shaded regions denote 95% confidence intervals. (**A**) Correlation between ESMRank score and baseline CFTR cell-surface abundance measured by immune-staining intensity (Spearman $\rho = 0.49$, $P = 2.63 \times 10^{-7}$). (**B**) Correlation between ESMRank score and response to elexacaftor (ELX) treatment ($\rho = 0.52$, $P = 2.5 \times 10^{-8}$). (**C**) Correlation between ESMRank score and response to tezacaftor (TEZ) treatment ($\rho = 0.51$, $P = 5.7 \times 10^{-8}$). (**D**) Correlation between ESMRank score and combined ELX + TEZ treatment response ($\rho = 0.54$, $P = 6.21 \times 10^{-9}$). Across all conditions, higher ESMRank scores—indicating lower predicted deleteriousness—were associated with increased baseline CFTR abundance and greater responsiveness to corrector treatment. These consistent positive relationships across independent experimental systems support a mechanistic link between predicted sequence constraint, folding efficiency, and pharmacological rescue capacity.

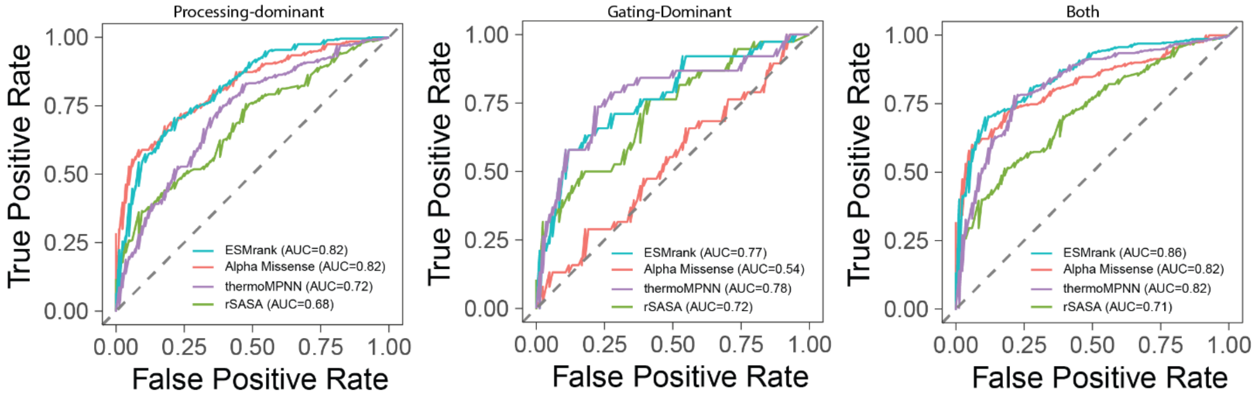


**Figure S12 | Mechanism-specific ROC analysis of CFTR pharmacological response classes.** Receiver operating characteristic (ROC) curves evaluating discrimination of mechanistic response subclasses defined in Fig. 4G. Variants were classified as processing-dominant (left), gating-dominant (middle), or responsive to both modulators (right) based on the relative contributions of corrector (ELX + TEZ) and potentiator (IVA) effects. In the processing-dominant class, ESMRank (AUC = 0.82) performed comparably to AlphaMissense (AUC = 0.82) and outperformed ThermoMPNN (AUC = 0.72) and solvent accessibility (rSASA; AUC = 0.68). In the gating-dominant class, ESMRank (AUC = 0.77) and ThermoMPNN (AUC = 0.78) retained strong discriminatory power, whereas AlphaMissense showed markedly reduced sensitivity (AUC = 0.54). For variants responsive to both modulators, ESMRank achieved the highest performance (AUC = 0.86), exceeding AlphaMissense (AUC = 0.82), ThermoMPNN (AUC = 0.82), and rSASA (AUC = 0.71). Across mechanistic subclasses, ESMRank maintained the highest mean discrimination (mean AUC = 0.82), outperforming ThermoMPNN (mean AUC = 0.77) and AlphaMissense (mean AUC = 0.73). This consistent performance across distinct therapeutic mechanisms supports a stability-centered continuum underlying both folding correction and gating potentiation.
